# Learning to Integrate an Artificial Sensory Device: Early Bayesian Integration and Conscious Perception

**DOI:** 10.1101/2020.01.29.924662

**Authors:** Mohammad-Ali Nikouei Mahani, Karin Maria Bausenhart, Rolf Ulrich, Majid Nili Ahmadabadi

**Affiliations:** Cognitive Systems Lab, School of Electrical and Computer Engineering, College of Engineering, University of Tehran, Tehran, Iran; Cognition and Perception, Department of Psychology, University of Tübingen, Tübingen, Germany

## Abstract

The present study examines how artificial tactile stimulation from a novel non-invasive sensory device is learned and integrated with information from another sensory system. Participants were trained to identify the direction of visual dot motion stimuli with a low, medium, and high signal-to-noise ratio. In bimodal trials, this visual direction information was paired with reliable symbolic tactile information. Over several training blocks, discrimination performance in unimodal tactile test trials and subjects’ confidence in their decision improved, indicating that participants were able to associate the visual and tactile information consciously and thus learned the meaning of the symbolic tactile cues. Formal analysis of the results in bimodal trials showed that both modalities are being integrated already in the early learning phases. Our modeling results revealed that this integration is consistent with a Bayesian model, which is an optimal integration of sensory information. Furthermore, we showed that a confidence-based Bayesian integration explains the observed behavioral data better than the classical variance-based Bayesian integration. Thus, the present study demonstrates that humans can consciously learn and integrate an artificial sensory device that delivers symbolic tactile information. This finding connects the field of multisensory integration research to the development of sensory substitution systems.

## 1. Introduction

Humans perceive their environment through multiple sensory inputs. Consequently, lacking or unreliable sensory inputs can cause a significant drop in the accuracy of perception. However, some artificial sensory devices, such as substitution systems, can partially compensate for the loss of a sensory modality (Abboud, Hanassy, Levy-Tzedek, Maidenbaum, & Amedi, 2014; Maidenbaum, Abboud, & Amedi, 2014), while others are designed to improve perception by providing complementary or processed information (Shull & Damian, 2015; Spiers & Dollar, 2016). For example, invasive artificial sensory systems exert a direct effect on the neuronal system (Collins et al., 2017) that can even lead to optimal integration of multisensory information (Dadarlat, O’Doherty, & Sabes, 2015). However, invasive techniques can still gain from further development. Therefore, non-invasive sensory feedback devices such as wearable systems are potentially the best alternatives for invasive techniques because they are already developed for realistic applications. Although there is a large number of studies that addressed the technical aspects of non-invasive and wearable devices (Iqbal, Aydin, Brunckhorst, Dasgupta, & Ahmed, 2016; Mukhopadhyay, 2015; Son et al., 2014), the cognitive aspects of them are less well studied. In particular, it is still unclear whether and how the input from a non-invasive artificial sensory device can be integrated into the multisensory perceptual system.

Many studies reported that adult humans integrate multiple sensory modalities in an optimal Bayesian fashion. Bayesian integration, in comparison to unimodal perception, leads to a significant decrease in response time (Diederich & Colonius, 2004; Drugowitsch, DeAngelis, Angelaki, & Pouget, 2015; Drugowitsch, DeAngelis, Klier, Angelaki, & Pouget, 2014) and an increase in accuracy and reliability of perception (Butler, Smith, Campos, & Bülthoff, 2010; Drugowitsch et al., 2015; Ernst & Banks, 2002; Mahani, Sheybani, Bausenhart, Ulrich, & Ahmadabadi, 2017; Pouget, Beck, Ma, & Latham, 2013). However, most of the previous studies have only examined the integration of well-experienced sensory inputs. Only a few studies addressed whether the learning of novel sensory devices leads to optimal integration or the selection of a sensory modality. Dadarlat et al. showed that monkeys could optimally integrate unfamiliar multichannel intracortical microstimulation (ICMS) signals and proprioceptive input(Dadarlat et al., 2015). However, so far, this issue has not been addressed for non-invasive artificial devices.

In the present study, we thus investigated the integration of visual motion information with symbolic input from an unfamiliar wearable vibrotactile device. Participants received visual random dot motion stimuli paired with synchronous static vibrotactile spatial patterns delivered through a vibrating belt in several consecutive training phases. They were asked to learn these visual-tactile associations. Following each training phase, the trained stimuli were presented either unimodally or bimodally and participants were asked to report the associated direction of motion. Accuracy and self-reported confidence were assessed to examine whether or not participants can learn the symbolic meaning conveyed by the wearable device and whether the information from the two inputs could be integrated. We also examined whether a similar change in confidence reports would accompany a variation in perceptual accuracy. According to previous studies(Overgaard, 2015; Rahnev, Denison, & Sciences, 2018; Vlassova, Donkin, & Pearson, 2014) consistent variation of perceptual performance and confidence reports is indicative of conscious perception.

In our experiment, participants performed a multisensory learning task that involves a novel artificial vibro-tactile device (see Figure 1**Error! Reference source not found.**). The experiment consisted of seven blocks, each involving a training phase followed by an evaluation phase. During the training phase, participants simultaneously received a dot motion pattern and a novel vibro-tactile stimulation pattern, which did not involve any directional movement. Throughout the whole experiment, each motion direction was paired with a specific symbolic vibro-tactile pattern. Participants were asked to learn the associations between each specific motion direction and the corresponding vibro-tactile stimulation pattern. Participants received unimodal visual motion dot stimuli, unimodal vibro-tactile stimuli, or multimodal stimuli combining both inputs (see materials and methods for more details) in the evaluation phase. Participants were then asked to report motion direction, either directly from the motion of the dots, from the associated vibro-tactile pattern, or from the combined multimodal stimulation. They also rated their confidence in these decisions. The visual stimuli in the unimodal and the multimodal condition had three levels of reliability: low, medium, and high, manipulated through the coherence of motion direction within the moving dot pattern. The reliability of tactile stimuli was constant during the whole experiment.

**Figure 1.**
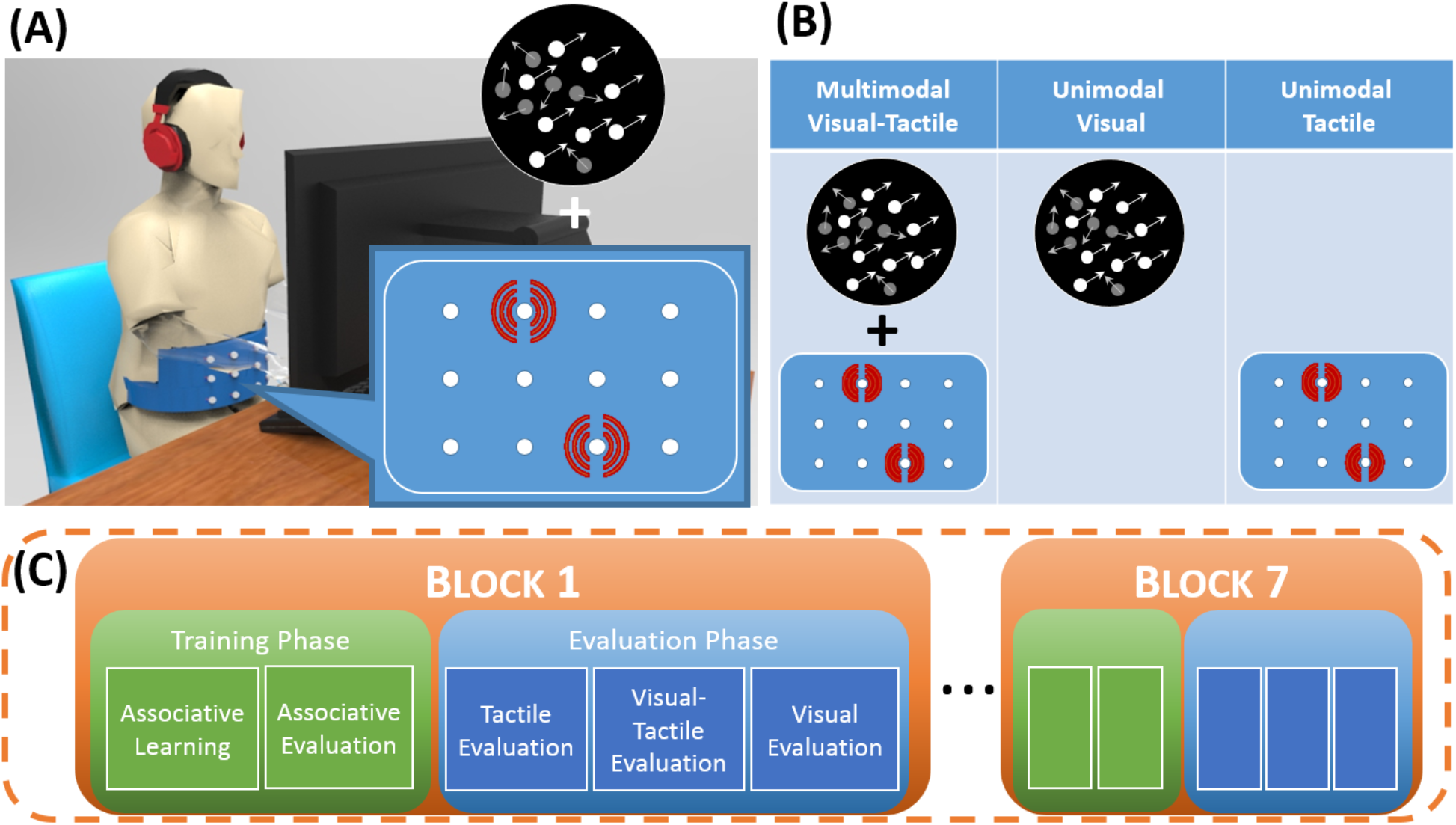
(A) Experimental setup. Participants received synchronous static vibrotactile spatial patterns and visual random dot motion stimuli. Vibroractile patterns were provided by a custom designed belt through a matrix of 3×4 tiny vibration motors. (B) Visual stimulus, tactile stimulus and synchronous visual-tactile stimulus. The reliability of visual dot motion stimulus was manipulated through three coherence levels, while the reliability of the vibrotactile stimulus was fixed. (C) The experiment consisted of seven blocks, each involving a training phase followed by an evaluation phase.

The impact of artificial sensory stimulation on multisensory perception was assessed over the course of learning. We analyzed the accuracy of unimodal and bimodal perception as well as confidence in the perceptual decisions from the evaluation phases of each of the seven consecutive blocks. For visual and bimodal trials, data were analyzed according to the low, medium, and high reliability of the visual dot motion stimuli.

## 2 Results

Figure 2 depicts the accuracy, reaction time, and confidence of perception in low, medium, and high reliability conditions across blocks. The accuracy of tactile perception increased over the course of the experiment, thus demonstrating that the visuo-tactile patterns provided by our artificial sensory device were efficiently learned and associated with the corresponding visual dot-motion patterns.

**Figure 2.**
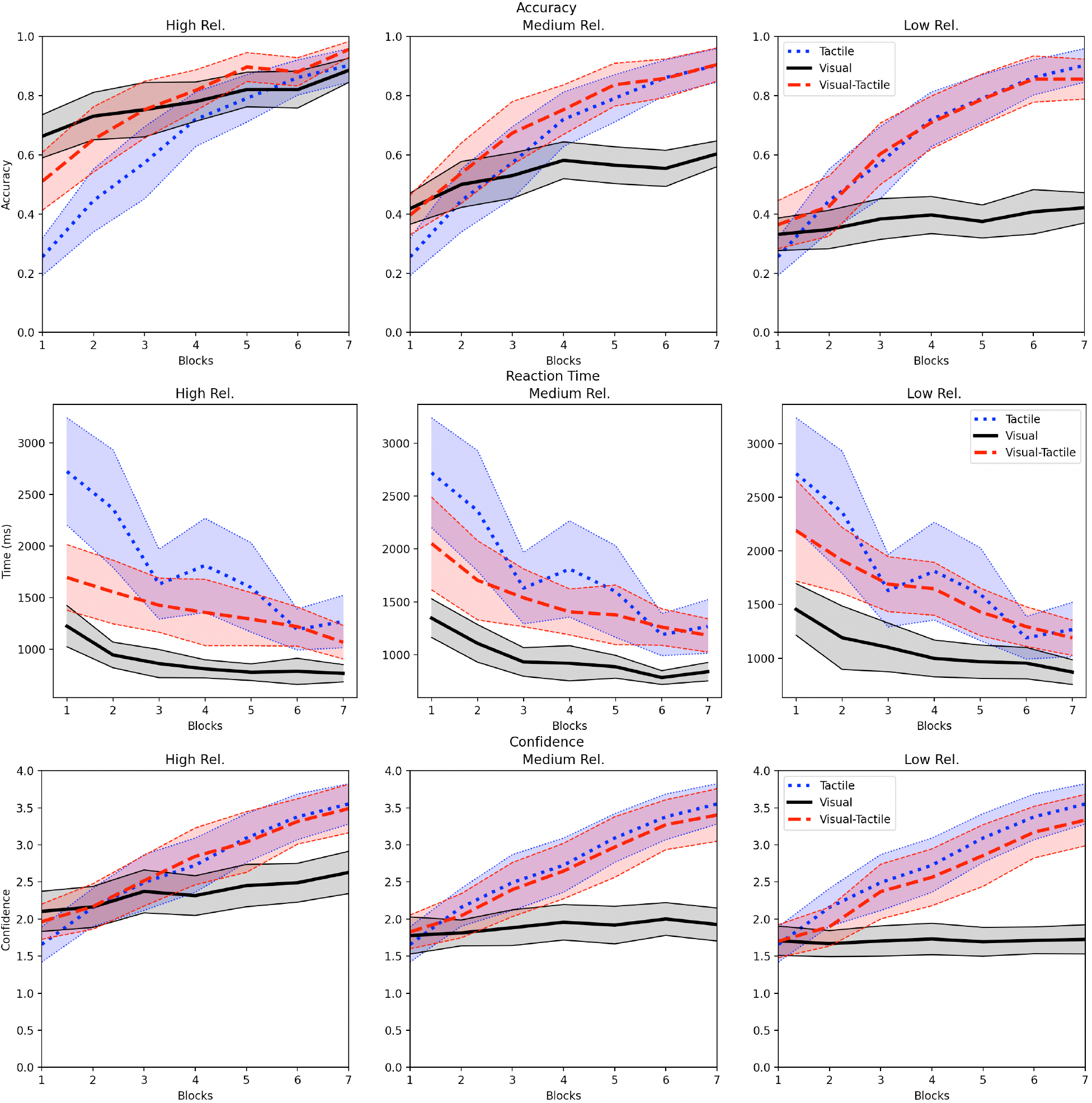
Accuracy, reaction time, and confidence of perception in low, medium, and high reliability conditions over the course of the learning procedure. For the sake of clarity, data from the tactile condition are re-plotted in each panel. Confidence intervals are corrected based on (Morey, 2008).

The experimental results show the integration of visual and tactile information in the accuracy and reaction time of perception. Integration behavior is specifically evident in the high-reliability condition, where the accuracy and reaction time of visual-tactile modality lay between unimodal visual and unimodal tactile from the early blocks. In the medium reliability condition, a similar pattern was evident regarding reaction time, while accuracy in visual-tactile trials surpassed unimodal accuracy in several blocks, suggesting integration of visual and tactile information in the middle of learning, but not in early and late blocks. In the low reliability condition, accuracy and, to some extent, reaction time for the visual-tactile trials was similar to unimodal tactile trials. This suggests that performance in bimodal trials is rather based on a selection of tactile information rather than an integration of tactile and visual information in this condition.

Confidence reports show that participants were aware of their perceptual performance, with confidence ratings reflecting the objective accuracy of perception. In all three reliability conditions, the confidence in tactile and visual-tactile modalities increased over the blocks of learning, consistent with the observed increase in perceptual accuracy. For the visual modality, confidence ratings showed a slight increase across learning blocks in the high reliability condition but did not change much in the low and medium reliability conditions, which is consistent with the pattern observed in perceptual accuracy.

Separate two-way within-subject ANOVAs with factors reliability condition (low, medium, and high) and block (block 1 to block 7) were performed on accuracy, confidence, and reaction time of visual and visual-tactile perceptions. The ANOVA analyses for confidence and reaction time are available in the supplementary material, while the ANOVA result for accuracy is described below.

The accuracy of unimodal visual perception was significantly affected by reliability condition, *F*(2,44) = 457.22, *p* < .001, as well as by block, *F*(6,132) = 9.48, *p* < .001. The interaction of reliability condition and block was not significant, *F*(12,264) = 1.47, *p* = .13. Post-hoc Tukey tests on reliability condition showed significant differences between all reliability conditions (*p*s < .001). This result points to a successful reliability control of the visual motion stimuli.

The analysis of accuracy in visual-tactile trials also showed a significant effect of reliability condition, *F*(2,44) = 30.73, *p* < .001. The effect of block on accuracy was significant, too, *F*(6,132) = 44.93, *p* < .001. Post-hoc Tukey tests on reliability condition showed the same result as for unimodal visual trials, that is, accuracy increased from low to medium, and from medium to high reliability (all *ps* < .001).

This effect of reliability condition was especially pronounced in the initial blocks of learning, as indicated by an interaction of both factors, *F*(12,264) = 2.81, *p* = .001. This shows that the combined information from both modalities is perceived differently depending on the reliability of the visual input.

Additional two-way within-subject ANOVAs with factors modality (visual, tactile, visual-tactile) and block (1 to 7) on accuracy, reaction time, and confidence were conducted for each of the three reliability conditions to assess learning and multisensory integration of the artificial sensory device. The ANOVA results for confidence and reaction time are available in the supplementary material.

In the low reliability condition, the effect of modality on accuracy, *F*(2,44) = 83.40, *p* < .001, the effect of block on accuracy, *F*(6,132) = 52.76, *p* < .001, and their interaction, *F*(12,264) = 23.04, *p* < .001, were significant. Post-hoc Tukey tests revealed that the visual stimuli were perceived less accurately than tactile (*p* < .001) and visual-tactile stimuli (*p* < .001). However, the difference between the accuracy of visual-tactile and tactile was not significant (*p* = .81). This result indicates a dominance of tactile stimulation on visual-tactile perception, when the reliability of the visual stimuli is low.

The same analysis in the medium reliability condition also showed effects of modality, *F*(2,44) = 24.93, *p* < .001, block, *F*(6,132) = 52.42, *p* < .001, and their interaction, *F*(12,264) = 16.08, *p* < .001. Although post-hoc Tukey tests indicated significant differences among all modalities (all *p*s < .007), the slope of accuracy across blocks was different between modalities. At the beginning of learning, the accuracy for the tactile stimuli was lower than for the visual stimuli, and the visual input determined the accuracy for visual-tactile stimulation. However, tactile perception got more accurate throughout learning, and as a result, its influence on the multimodal visual-tactile perception increased. In some learning blocks, the accuracy for visual-tactile stimulation was even higher than for the unimodal visual and tactile stimuli in isolation (see Figure 2). This indicates an effective integration of visual and tactile information (see the modeling section for a more elaborated account of this integration effect).

Finally, in the high reliability condition, accuracy was also affected by modality, *F*(2,44) = 13.59, *p* < .001, block, *F*(6,132) = 47.80, *p* < .001, and their interaction, *F*(12,264) = 14.67, *p* < .001. At the beginning of the experiment, the accuracy for visual-tactile stimulation was codetermined by both visual and tactile information, as indicated by intermediate accuracy, lying between the respective accuracies for visual and tactile unimodal stimulation. This was confirmed by a one-way within-subject ANOVA, which showed a significant difference of accuracy between the different modalities in the first block, *F*(2,44) = 55.21, *p* < .001. Post-hoc tests showed significant differences among all modalities (*p*s < .001). Most interestingly, this suggests that participants integrated the visual and tactile information from the beginning of learning, even though this impaired performance compared to the unimodal visual condition.

### Evidence for conscious integration

As already mentioned, the change of confidence reports is consistent with the change of perceptual accuracy in all three conditions. Specifically, we can see a bold increase in confidence reports for tactile and visual-tactile modalities across blocks, which is consistent with the significant increase of accuracy in these modalities, see Figure 2. However, in the visual modality, the increase of confidence and accuracy is much less prominent, presumably due to familiarity with the stimulation and, therefore, lower perceptual learning in the visual modality.

As Figure 3 depicts, the correlation between confidence reports and accuracy in the tactile and the visual-tactile modality is higher than in the visual modality. Tactile perception started with low accuracy and low confidence reports, however, both accuracy and confidence reports increased throughout blocks. This shows that participants were aware of their own performance during the learning of the novel tactile device and could successfully adjust their confidence reports. A similar pattern can be observed for visual-tactile stimulation in all reliability conditions. Although for the unimodal visual stimuli, changes in accuracy and confidence reports were less prominent, the correlation between accuracy and confidence reports was still positive and significant. In summary, we can conclude that participants are aware of their learning and multimodal perception of a novel sensory device, and both the novel and familiar stimulations are accessible to conscious processing.

**Figure 3.**
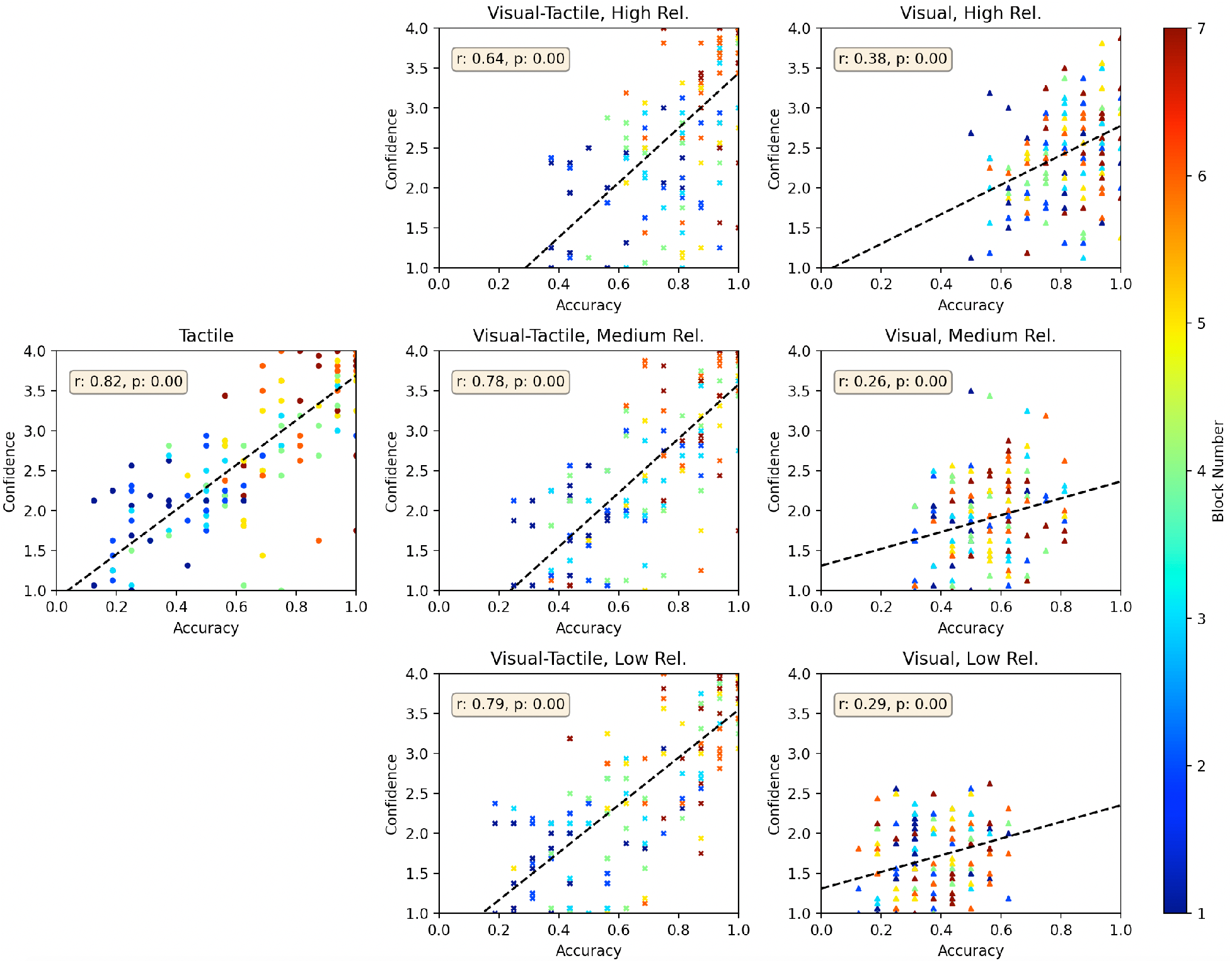
Correlation of confidence reports and accuracy in all reliability conditions and for all modalities. The color map shows the block numbers. Dashed black lines show the fitted linear regression line and *r* and *p* value of Pearson correlation are indicated for each condition. For accuracy, outlier values (outside the mean ± 2 σ range) were excluded from the plots and analyses.

## 3 Modeling

The potential mechanisms underlying the integration of the two modalities were investigated by computational modeling. In order to shed light on the mechanisms behind the perceptual learning of this artificial sensory device, we compared the Bayesian integration model to the selection model for both first-order (perceptual decisions) and second-order (confidence reports) decisions.

### 3.1 Confidence Confusion Matrix; Second Order Confusion Matrix

The confusion matrix (CM) is a valuable tool for evaluating the performance of participants in a multiple-choice task or the performance of a classifier algorithm in solving supervised multi-class problems. Various criteria like accuracy, precision, and recall can be determined from the CM for both 2AFC and multiple choice (multiple class) problems.

For the present purpose, the confusion matrix is extended to second-order judgments, that is, self-reported confidence in judgments. When a participant (a classifier) reports confidence for each of the first-order judgments, this meta-information can be used to compute meta-accuracy (type II accuracy). A confidence confusion matrix (*CCM*) is defined with the same structure as the confusion matrix as follows:

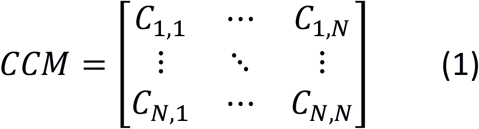

*C*_*i,j*_ indicates the sum of all confidence ratings associated with response *R*_*i,j*_, where a ground truth signal *i* (class *i*) has been predicted as signal *j* (class *j*). Similar to the accuracy, we define meta-accuracy as the sum of the diagonal divided by the sum of all elements:

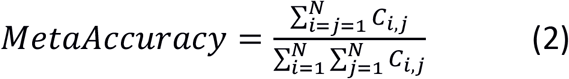

Meta-accuracy represents the performance of the participants not only by considering their decisions but also by weighting them according to the confidence ratings associated with these decisions. Thus, high-confidence correct answers increase meta-accuracy more than low-confidence correct answers, while the low-confidence wrong answers decrease the meta-accuracy less than the high confidence wrong answers. In fact, CCM is a more generic representation of performance than CM, which is a special case with just one confidence level.

### 3.2 Bayesian Integration vs Selection Model

During the previous decade, multisensory integration has become an increasingly prominent research topic and various computational models have been proposed to unravel the mechanism underlying sensory integration across multiple modalities (Diederich & Colonius, 2004; Drugowitsch et al., 2015; Drugowitsch et al., 2014; Mahani et al., 2017). The most prominent model of multisensory integration suggests that the sensory inputs from different modalities are combined according to the principle of maximum-likelihood estimation (MLE)(Ernst & Banks, 2002). Specifically, this model holds that the inputs from different modalities are linearly combined according to the reliability of their respective sensory inputs. This simply means that if there are two modalities, A and B, the weight of each modality in integration, *w_A_*, and *w_B_* = 1 − *w_A_*, and the optimal integrated signal *S_AB_* are calculated as follows:

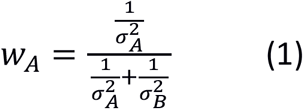

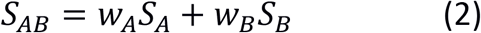

where 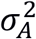 and 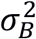 show the variance of signal A and variance of signal B, respectively.

In the following, we investigate how well this Bayesian model can predict the visual-tactile perception observed in our experiment based on the observed variance of unimodal visual and tactile perception.

Each row of a CM/CCM represents a true signal and columns show the decisions/predictions. Thus, we can estimate the variance of each true signal, *i*, by estimating the variance of all possible decisions, *j*, of the corresponding row. We defined the variance in perception as follows:

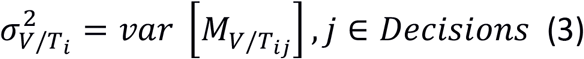

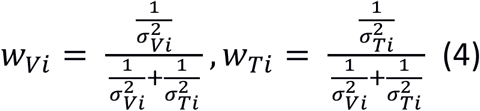

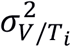 is the variance of true signal *i*, for the modality *V/T* based on the Matrix M. Matrix M is either CM or CCM in our modelling. Specifically, we calculated the variance of *i*^th^ row of the CM/CCM for modality *V/T*. *w_Vi_* and *w_Ti_* denote the weight of visual and tactile modalities, respectively.

In the present study, we compared the Bayesian integration model to the selection model for the first and the second-order judgments. As outlined above, in the Bayesian integration model, the information from two modalities is integrated according to the reliability of the unimodal contributions. In contrast, in a selection model, the information of the most reliable sensor is selected for each signal (each row of CM/CCM). To compare these two models, we fitted a Bayesian and a selection model to the first order (CM) and second order (CCM) observed experimental data. We therefore defined the predicted CM/CCM by the Bayesian/selection model as follows:

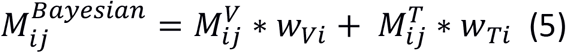

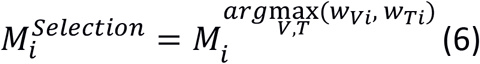

*M^Bayesian^* shows the predicted CM/CCM by the Bayesian integration model, and *M^Selection^* denotes the predicted CM/CCM by the selection model. In the Bayesian model, each row of the predicted CM/CCM is the weighted average of both sensory inputs for the corresponding row. However, in the selection model, each row of the predicted CM/CCM is selected, as a whole, from the most reliable sensor. Since each row of the CM/CCM represents the decision distribution of a specific signal, the weighted average in the Bayesian model would integrate the decision distributions from both modalities. In contrast, the selection model selects the most reliable decision distribution for each specific signal among the sensory modalities. By increasing the difference between the variance of two modalities, the Bayesian Integration model resembles more and more the selection model. When a modality has significantly higher variance than the other modality, its weight (in Eq. 5) becomes very small and neglectable. Thus, the Bayesian integration model resembles the selection model.

Figure 4 depicts the predicted accuracy by the Bayesian integration and the selection model based on first order (CM) and second-order (CCM) judgments. The Mean Absolute Error (MAE) between the observed experimental data and the model prediction is plotted for each condition. The second-order Bayesian integration model achieved the minimum MAE in all reliability conditions and thus, describes the behavior of multisensory perception better than the other models.

**Figure 4.**
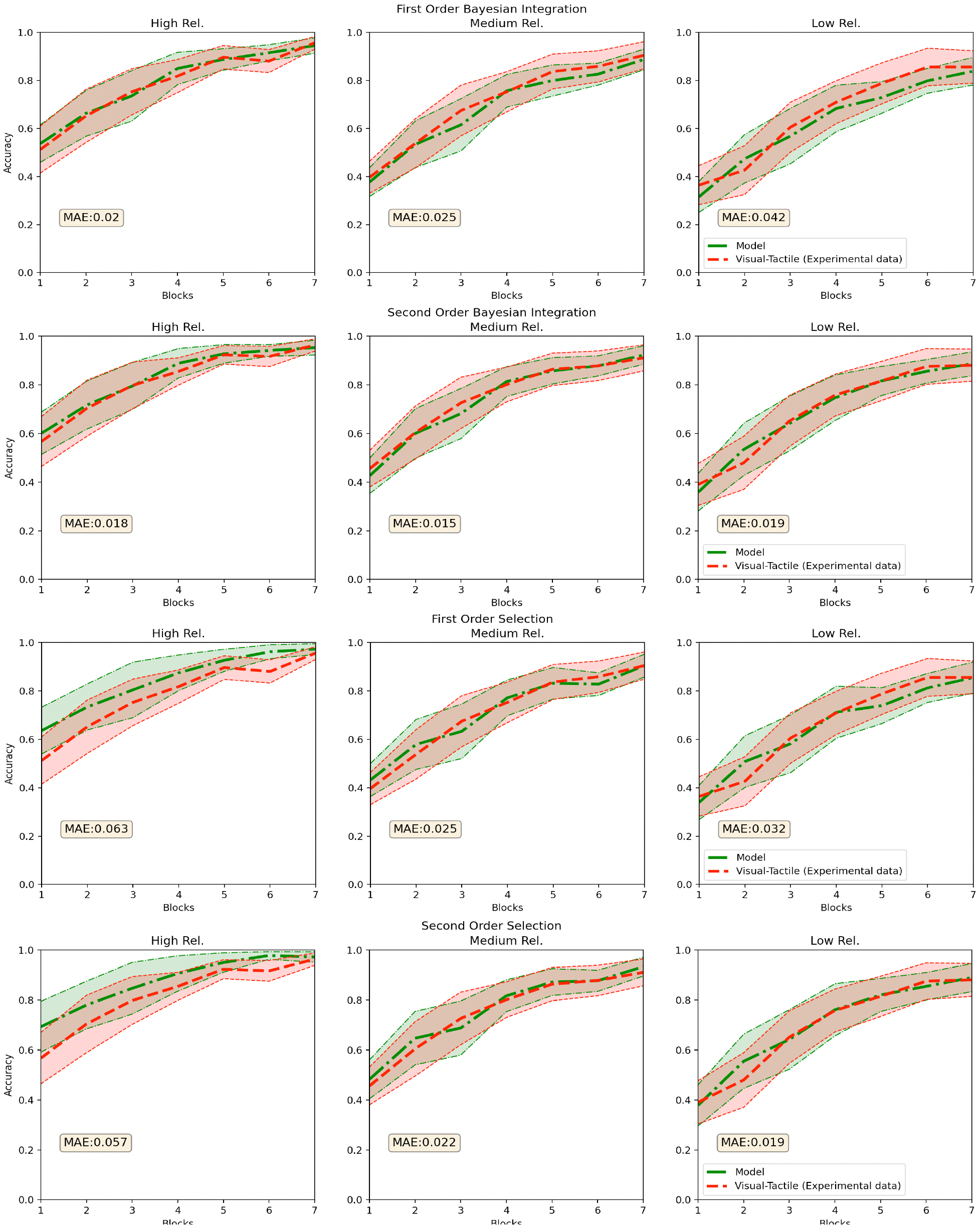
Observed and modeled accuracies in the visual-tactile condition. Data were modeled based on the first/second-order judgments observed in the unimodal conditions according to a Bayesian integration (Rows 1 and 2) and a selection model (Rows 3 and 4). Models were fitted to each subject, and the confidence intervals were calculated based on (Morey, 2008).

In the high-reliability condition, both Bayesian models fit the observed data better than both selection models. Both selection models systematically overestimate the observed accuracy and thus fail to capture perception in visual-tactile trials. In the medium reliability condition, all models could fairly fit the observed experimental data. Nevertheless, the second-order Bayesian integration model achieved the smallest MAE. In the low-reliability condition, the observed accuracy for the visual-tactile condition closely follows the observed unimodal tactile accuracy, suggesting selection behavior rather than integration. Thus, it explains why the first-order Bayesian integration model could not reasonably predict the perception behavior, and in contrast, both selection models fit the observed data better than the first-order Bayesian integration model. However, the reasonable fit of the second-order Bayesian integration model supports the generalization of this model also to the low-reliability condition. Taking all results into account, the second-order Bayesian integration model seems to be a universal model that can reasonably describe multisensory perception in all reliability conditions.

## Discussion

With the advance of medical and wearable stimulation devices, understanding the mechanisms of perception for such novel artificial devices is of great interest to scientists and engineers. This study addresses whether and how symbolic information from a novel non-invasive artificial sensory device is learned and becomes integrated within the human sensory system. We have introduced a custom-designed vibrotactile belt that generates novel and unexperienced tactile patterns. Participants were asked to learn the associations between these novel tactile stimuli and the direction of visual dot motion stimuli across seven consecutive training phases. At the end of each training phase, there was an evaluation phase in which the performance in unimodal visual, unimodal tactile, and visual-tactile conditions was assessed. Thus, perceptual accuracy for the different modality conditions could be tracked over the time-course of the whole experiment.

The results show that accuracy in the tactile condition increased throughout the experiment, and thus, participants could learn the novel tactile patterns. The results also revealed that symbolic information from the novel artificial device can be integrated into the visual-tactile perception from the very beginning of the training. This integration process was evident in the high visual reliability condition, even though accuracy for the tactile modality was initially lower than for the visual modality. In the low visual reliability condition, the tactile information dominated over the visual information and seemingly resulting in selection behavior rather than sensory integration. However, as already mentioned in the modeling section, when the difference between variance of two modalities is large, the Bayesian integration model might resemble the selection model and that could be the case in the low reliability condition. The modeling results also equally support the Bayesian integration and selection models in the low-reliability condition.

The assessment of confidence reports showed that participants’ confidence reports consistently reflected their accuracy during the learning of the novel tactile device: accuracy, as well as confidence in tactile perception, increased over the course of learning, and both measures correlated positively and significantly. The consistent alteration of accuracy and confidence shows that participants were aware of their own perception during the learning and perception of the novel tactile sensory device [24, 25]. Similar results were observed for visual-tactile stimulation, where both accuracy and confidence values also increased over the course of learning. The results regarding confidence reports thus provide some evidence for conscious learning and integration of the novel tactile device, which is an important finding for developing and utilizing such artificial devices.

To better understand the mechanisms underlying the perception and integration of novel symbolic information, the predictions of four computational models were compared with respect to the visual-tactile data of this study: Bayesian integration of first/second-order judgments and selection based on the first/second-order judgments. The Bayesian integration model assumes an optimal integration of sensory information based on the variance of the first/second-order judgments for unimodal inputs. In contrast, the selection model assumes that the information from the unimodal sensor with the lowest variance in first/second-order judgments is selected. The modeling results show that both the first and the second-order Bayesian integration model could fit the experimental data marginally better than the selection models in the high-reliability condition, where the integration behavior was also prominant in the observed experimental data. In the medium reliability condition, all models could fairly fit the observed experimental data, but the second-order Bayesian model fitted slightly better than the others. In the low-reliability condition, where the observed data seemingly reflects selection behavior, the first-order Bayesian integration model fits the observed data worse than both selection models and the second-order Bayesian model. Taking all together, the second-order Bayesian integration model achieved the minimum MAE across reliability conditions and thus seems like a generic model that allows for the most parsimonious account of the data. Since CCM is a richer and more generic representation of the performance than CM, it allows the second-order Bayesian model to precisely fit the observed data in all reliability conditions. Our findings are consistent with other visual-tactile studies with experienced/natural sensory information, showing that the information from visual and tactile modalities is integrated in an Optimal Bayesian fashion (Burge, Girshick, & Banks, 2010).

In conclusion, the present data show that participants can consciously utilize symbolic tactile information to improve processing of visual-tactile information, thereby improving the accuracy of perception even in the early phases of learning. Our modeling results showed that the Bayesian model with access to the second-order judgments explains visual-tactile perception better than a Bayesian model with access to first-order judgments and selection models with access to first or second-order judgments. Accordingly, the present study revealed that symbolic information from a novel tactile device optimally integrates with the visual information from the early stages of learning. Although many studies showed the optimal Bayesian integration of visual and tactile information, some studies provided evidence for non-Bayesian integration. (Adams, 2019; van Beers, van Mierlo, Smeets, & Brenner, 2011). We believe the present paper can provide insights for developing further models of multisensory integration based on second-order judgments. Second-order models can potentially reduce the gap between the Bayesian and non-Bayesian models.

## 5. Methods

### Participants

18 women and 11 men (23.72 ± 3.42 years old) recruited from a student population of Tübingen University participated in the experiment. They all reported normal or corrected-to-normal vision and no neurological or psychiatric disorder. The experiment was conducted under the Helsinki Declaration and the guidelines for scientific work of the University of Tübingen. Written informed consent was obtained from all participants before data collection. Participants were compensated with 8 Euros per hour or course credit for their participation. They were also told that the best performer would get a 50 Euro Amazon gift card. The results of six participants were excluded from all analyses, since their performance in tactile perception was always below 50% in all blocks.

### Stimuli and Apparatus

Participants were seated in a sound-attenuated room in front of a 19” screen with 100 Hz refresh rate and 1024×768 pixels resolution, on which the visual stimuli were presented. The distance between participants’ eyes and the monitor screen was about 50 cm. The experiment was implemented in C++.

The visual stimuli consisted of a random dot motion display, made up of 100 white dots against a black background. The diameter of each dot was 10 pixels, approximately 0.33° visual angle. The dots were randomly distributed within a circle with a diameter of 400 pixels (approximately 13.3° visual angle) called excircle. The initial x/y position of each dot came from two Gaussian distributions with a mean zero (center of the screen) and variances of 0.1 of the width/height of the screen. Each dot started to move with a speed of 11.25°/s in a specific direction. Whenever a dot hit the excircle, it was replaced by a new dot. There were eight possible directions (0, 45, 90, 135, 180, 225, 270, and 315°). The percentage of the dots moving in the same direction, called coherence level, determined the reliability of the visual stimulus. For example, a coherence level of 75% means 75 out of 100 dots moved in the same direction, while 25 dots moved in a random direction. The duration of the visual stimulus was one second.

Tactile stimuli were provided by a custom-designed belt fastened around the subjects’ waist. The vibrotactile belt was designed in the Cognitive Robotics Lab at University of Tehran. It consists of 12 vibration motors located in a 3×4 matrix formation on a cotton canvas tape. All motors are controlled by a custom embedded device similar to the one employed in a previous study. (Mahani et al., 2017) Eight different tactile stimulation patterns were produced, each consisting of two simultaneous vibrations at pre-defined locations on the belt. These patterns were defined to be maximally distinguishable. The vibration of all tactile stimuli lasted one second, that is, the same duration as the visual stimuli. White-noise auditory stimuli were continuously delivered to the subjects through a pair of Sony MDR-XD200 stereo headphones to override the sound produced by the vibration motors.

### Procedure

The experiment consisted of seven consecutive blocks, each of which was divided into a *training phase* and an *evaluation phase (*see Figure 1**Error! Reference source not found.**).

#### Training phase

Each block started with a training phase in which participants were asked to learn the associations between the tactile patterns and the visual motion directions. A random, subject-specific, and one-by-one mapping associates each tactile pattern with one of the visual motion directions. The mappings were randomly assigned before the experiment for each participant. The training phase included an associative learning section followed by an associative test section. In the learning section, all visual-tactile association pairs were presented four times in random order. In the associative test section, participants were asked to decide if the presented pair of visual-tactile stimuli were associated or not by selecting a green/red circle with a mouse click. Subsequently, they reported their confidence in these decisions on a scale of 1 (lowest confidence level) to 4 (highest confidence level) by selecting one of four adjacent circles by mouse. The cursor position was set to the screen center before each judgment to avoid response bias. Participants received feedback about the correctness of their decision at the end of each trial. A splash screen, which showed the overall accuracy achieved during the test section, concluded each training phase. The coherence of the visual stimuli during the whole training phase was 75 %.

#### Evaluation phase

Following each training phase, performance for visual, tactile, and visual-tactile judgments was assessed. Each evaluation phase included a visual evaluation section, a tactile evaluation section, and a visual-tactile evaluation section, which were presented in random order. In each trial of the visual evaluation section, participants received a unimodal visual stimulus with a coherence level of 10%, 15%, or 25% (low, medium, and high reliability, respectively). Participants were asked to judge the direction of the dot motion by a mouse click on the corresponding sector in a circle. Then, they were asked to report their confidence level on a scale of 1 to 4, similar to the test section of the training blocks. The visual evaluation section consisted of 48 trials (8 directions x 3 coherence levels x 2 repetitions), presented in random order. The visual-tactile evaluation section was identical to the unimodal visual evaluation section, except that its associated tactile pattern accompanied the visual stimulus in each trial. The tactile evaluation section consisted of 16 trials (8 patterns x 2 repetitions). Participants were asked to report the learned associated motion direction and their confidence in this decision. In this phase, no feedback was provided.

