## Supplementary for "Learning to Integrate an Artificial Sensory Device: Early Bayesian Integration and Conscious Perception"

### ANOVA Analysis

#### ANOVA on Confidence with factors block and reliability

##### Visual Modality

|  | SS | ddof1 | ddof2 | MS | F | p-unc | p-GG-corr | np2 | eps |
| --- | --- | --- | --- | --- | --- | --- | --- | --- | --- |
| Reliability | 36.49 | 2 | 44 | 18.24 | 82.51 | 0.0 | 0.00 | 0.79 | 0.60 |
| Block | 3.16 | 6 | 132 | 0.53 | 3.33 | 0.0 | 0.02 | 0.13 | 0.54 |
| Reliability * Block | 2.44 | 12 | 264 | 0.20 | 5.54 | 0.0 | 0.00 | 0.17 | 0.30 |

##### Visual-Tactile Modality

|  | SS | ddof1 | ddof2 | MS | F | p-unc | p-GG-corr | np2 | eps |
| --- | --- | --- | --- | --- | --- | --- | --- | --- | --- |
| Reliability | 3.44 | 2 | 44 | 1.72 | 30.52 | 0.00 | 0.00 | 0.58 | 0.77 |
| Block | 146.97 | 6 | 132 | 24.49 | 37.81 | 0.00 | 0.00 | 0.63 | 0.57 |
| Reliability * Block | 0.38 | 12 | 264 | 0.03 | 1.21 | 0.27 | 0.31 | 0.05 | 0.20 |

#### ANOVA on Confidence with factors block and modality

##### Low Reliability

|  | SS | ddof1 | ddof2 | MS | F | p-unc | p-GG-corr | np2 | eps |
| --- | --- | --- | --- | --- | --- | --- | --- | --- | --- |
| Modality | 95.70 | 2 | 44 | 47.85 | 82.97 | 0.0 | 0.0 | 0.79 | 0.61 |
| Block | 77.89 | 6 | 132 | 12.98 | 50.96 | 0.0 | 0.0 | 0.70 | 0.55 |
| Modality * Block | 37.96 | 12 | 264 | 3.16 | 24.69 | 0.0 | 0.0 | 0.53 | 0.23 |

##### Medium Reliability

|  | SS | ddof1 | ddof2 | MS | F | p-unc | p-GG-corr | np2 | eps |
| --- | --- | --- | --- | --- | --- | --- | --- | --- | --- |
| Modality | 67.42 | 2 | 44 | 33.71 | 68.73 | 0.0 | 0.0 | 0.76 | 0.63 |
| Block | 83.33 | 6 | 132 | 13.89 | 49.15 | 0.0 | 0.0 | 0.69 | 0.51 |
| Modality * Block | 30.81 | 12 | 264 | 2.57 | 21.67 | 0.0 | 0.0 | 0.50 | 0.25 |

##### High Reliability

|  | SS | ddof1 | ddof2 | MS | F | p-unc | p-GG-corr | np2 | eps |
| --- | --- | --- | --- | --- | --- | --- | --- | --- | --- |
| Modality | 15.79 | 2 | 44 | 7.90 | 13.46 | 0.0 | 0.0 | 0.38 | 0.60 |
| Block | 93.56 | 6 | 132 | 15.59 | 56.47 | 0.0 | 0.0 | 0.72 | 0.52 |
| Modality * Block | 19.82 | 12 | 264 | 1.65 | 12.79 | 0.0 | 0.0 | 0.37 | 0.26 |

#### ANOVA on Reaction Time with factors block and reliability

##### Visual Modality

|  | SS | ddof1 | ddof2 | MS | F | p-unc | p-GG-corr | np2 | eps |
| --- | --- | --- | --- | --- | --- | --- | --- | --- | --- |
| Reliability | 3096411.06 | 2 | 44 | 1548205.53 | 11.77 | 0.00 | 0.00 | 0.35 | 0.59 |
| Block | 13796588.00 | 6 | 132 | 2299431.33 | 22.19 | 0.00 | 0.00 | 0.50 | 0.35 |
| Reliability * Block | 358797.29 | 12 | 264 | 29899.77 | 0.89 | 0.56 | 0.43 | 0.04 | 0.19 |

##### Visual-Tactile Modality

|  | SS | ddof1 | ddof2 | MS | F | p-unc | p-GG-corr | np2 | eps |
| --- | --- | --- | --- | --- | --- | --- | --- | --- | --- |
| Reliability | 5027583.37 | 2 | 44 | 2513791.68 | 16.07 | 0.00 | 0.00 | 0.42 | 0.80 |
| Block | 33364420.14 | 6 | 132 | 5560736.69 | 11.36 | 0.00 | 0.00 | 0.34 | 0.45 |
| Reliability * Block | 1914175.49 | 12 | 264 | 159514.62 | 1.45 | 0.14 | 0.24 | 0.06 | 0.23 |

#### ANOVA on Reaction Time with factors block and modality

##### Low Reliability

|  | SS | ddof1 | ddof2 | MS | F | p-unc | p-GG-corr | np2 | eps |
| --- | --- | --- | --- | --- | --- | --- | --- | --- | --- |
| Modality | 45484457.77 | 2 | 44 | 22742228.89 | 40.14 | 0.0 | 0.00 | 0.65 | 0.73 |
| Block | 54998458.12 | 6 | 132 | 9166409.69 | 23.07 | 0.0 | 0.00 | 0.51 | 0.45 |
| Modality * Block | 10880172.61 | 12 | 264 | 906681.05 | 3.58 | 0.0 | 0.02 | 0.14 | 0.28 |

##### Medium Reliability

|  | SS | ddof1 | ddof2 | MS | F | p-unc | p-GG-corr | np2 | eps |
| --- | --- | --- | --- | --- | --- | --- | --- | --- | --- |
| Modality | 56112559.81 | 2 | 44 | 28056279.91 | 45.70 | 0.0 | 0.0 | 0.68 | 0.67 |
| Block | 49706609.30 | 6 | 132 | 8284434.88 | 17.91 | 0.0 | 0.0 | 0.45 | 0.48 |
| Modality * Block | 11083544.06 | 12 | 264 | 923628.67 | 4.84 | 0.0 | 0.0 | 0.18 | 0.24 |

##### High Reliability

|  | SS | ddof1 | ddof2 | MS | F | p-unc | p-GG-corr | np2 | eps |
| --- | --- | --- | --- | --- | --- | --- | --- | --- | --- |
| Modality | 67751676.12 | 2 | 44 | 33875838.06 | 43.65 | 0.0 | 0.0 | 0.66 | 0.82 |
| Block | 38220518.30 | 6 | 132 | 6370086.38 | 18.99 | 0.0 | 0.0 | 0.46 | 0.48 |
| Modality * Block | 14987420.65 | 12 | 264 | 1248951.72 | 6.12 | 0.0 | 0.0 | 0.22 | 0.23 |

#### ANOVA on Accuracy with factors block and reliability

##### Visual Modality

|  | SS | ddof1 | ddof2 | MS | F | p-unc | p-GG-corr | np2 | eps |
| --- | --- | --- | --- | --- | --- | --- | --- | --- | --- |
| Reliability | 13.0 | 2 | 44 | 6.50 | 457.22 | 0.00 | 0.0 | 0.95 | 0.96 |
| Block | 1.20 | 6 | 132 | 0.20 | 9.49 | 0.00 | 0.0 | 0.30 | 0.66 |
| Reliability * Block | 0.19 | 12 | 264 | 0.02 | 1.47 | 0.13 | 0.2 | 0.06 | 0.44 |

##### Visual-Tactile Modality

|  | SS | ddof1 | ddof2 | MS | F | p-unc | p-GG-corr | np2 | eps |
| --- | --- | --- | --- | --- | --- | --- | --- | --- | --- |
| Reliability | 1.24 | 2 | 44 | 0.62 | 30.74 | 0.0 | 0.00 | 0.58 | 0.66 |
| Block | 13.38 | 6 | 132 | 2.23 | 44.94 | 0.0 | 0.00 | 0.67 | 0.47 |
| Reliability * Block | 0.28 | 12 | 264 | 0.02 | 2.81 | 0.0 | 0.05 | 0.11 | 0.23 |

#### ANOVA on Accuracy with factors block and modality

##### Low Reliability

|  | SS | ddof1 | ddof2 | MS | F | p-unc | p-GG-corr | np2 | eps |
| --- | --- | --- | --- | --- | --- | --- | --- | --- | --- |
| Modality | 8.02 | 2 | 44 | 4.01 | 83.40 | 0.0 | 0.0 | 0.79 | 0.62 |
| Block | 9.97 | 6 | 132 | 1.66 | 52.77 | 0.0 | 0.0 | 0.71 | 0.58 |
| Modality * Block | 3.45 | 12 | 264 | 0.29 | 23.04 | 0.0 | 0.0 | 0.51 | 0.31 |

##### Medium Reliability

|  | SS | ddof1 | ddof2 | MS | F | p-unc | p-GG-corr | np2 | eps |
| --- | --- | --- | --- | --- | --- | --- | --- | --- | --- |
| Modality | 2.48 | 2 | 44 | 1.24 | 24.93 | 0.0 | 0.0 | 0.53 | 0.60 |
| Block | 10.63 | 6 | 132 | 1.77 | 52.42 | 0.0 | 0.0 | 0.70 | 0.49 |
| Modality * Block | 2.40 | 12 | 264 | 0.20 | 16.08 | 0.0 | 0.0 | 0.42 | 0.27 |

##### High Reliability

|  | SS | ddof1 | ddof2 | MS | F | p-unc | p-GG-corr | np2 | eps |
| --- | --- | --- | --- | --- | --- | --- | --- | --- | --- |
| Modality | 1.82 | 2 | 44 | 0.91 | 13.59 | 0.0 | 0.0 | 0.38 | 0.69 |
| Block | 9.83 | 6 | 132 | 1.64 | 47.80 | 0.0 | 0.0 | 0.68 | 0.43 |
| Modality * Block | 1.98 | 12 | 264 | 0.16 | 14.67 | 0.0 | 0.0 | 0.40 | 0.28 |
